# How do I bite thee? Let me count the ways: Exploring the Implications of Individual Biting Habits of Aedes aegypti for Dengue Transmission

**DOI:** 10.1101/2021.09.07.459140

**Authors:** Rebecca C. Christofferson, Helen J. Wearing, Erik A. Turner, Christine S. Walsh, Henrik Salje, Cécile Tran Kiem, Simon Cauchemez

## Abstract

In models of mosquito-borne transmission, the mosquito biting rate is an influential parameter, and understanding the heterogeneity of the process of biting is important, as biting is usually assumed to be relatively homogeneous across individuals, with time-between-bites described by an exponentially distributed process. However, these assumptions have not been addressed through laboratory experimentation. We experimentally investigated the daily biting habits of *Ae. aegypti* at three temperatures (24°C, 28°C, and 32°C) and determined that there was individual heterogeneity in biting habits (number of bites, timing of bites, etc.). We further explored the consequences of biting heterogeneity using an individual-based model designed to examine whether a particular biting profile determines whether a mosquito is more or less likely to 1) become exposed given a single index case of dengue (DENV) and 2) transmit to a susceptible human individual. Our experimental results indicate that there is heterogeneity among individuals and among temperature treatments. We further show that this results in altered probabilities of transmission of DENV to and from individual mosquitoes based on biting profiles. While current model representation of biting may work under some conditions, it might not uniformly be the best fit for this process. Our data also confirm that biting is a non-monotonic process with temperatures around 28°C being optimum.

**Author Summary:** Mosquito biting is a necessary and critical part of arbovirus transmission. The mosquito must bite once to acquire a virus and again to transmit, and these two bites must be separated by sufficient time for the virus to get to the salivary glands of the mosquito. Thus, both the number and timing of bites is important. We experimentally investigated how these bite characteristics might be different among individuals and further explored how temperature affected the overall heterogeneity of biting in *Aedes aegypti* mosquitoes, which carry many arboviruses like dengue virus (DENV). We found that the biting profiles – including number and timing thereof – did vary within temperature groups among individuals and compared outcomes associated with each individual in an individual based model of household DENV transmission. Our results further confirmed that temperatures around 28°C are optimal for mosquito biting (and transmission), that correlations between biting characteristics and transmission were not uniform across temperature, which represents another layer of heterogeneity, and that – at least at 28°C – the null assumption of an exponential or an exponential like (geometric) distribution of biting in mathematical models of transmission is not the best and offer an alternative.

## Introduction

*Aedes aegypti* are the primary vectors for several arboviruses of public health importance and are primarily found in tropical regions. *Ae. aegypti* tend to be in urban areas and often live close to or within human dwellings with typically limited flight ranges. There are usually abundant opportunities for daily blood meals from human hosts [1-4]. Bloodmeal analyses from mosquito trapping experiments have identified that the majority of bloodmeals in female *Ae. aegypti* mosquitoes are taken from humans that live in the same household where the mosquito was found [5]. Moreover, *Ae. aegypti* are known to take multiple bloodmeals – sometimes from multiple individuals – within a single gonotrophic cycle [5-7]. These bites provide opportunity for transmission of *Aedes*-borne viruses such as dengue (DENV), which is primarily moved by humans rather than mosquitoes among households [8]. The transmission system is therefore reliant upon contact between susceptible mosquito vectors and infectious and susceptible humans.

Biting is an influential component of the transmission cycle of *Ae. aegypti-*borne viruses [9-17]. Mosquitoes must bite once to acquire a virus and again to transmit that virus [18, 19]. The timing of these two events is also important, as these two bites must be separated by a sufficient period such that the virus can disseminate through the mosquito and establish an infection in the salivary glands [18]. After this period of time, called the extrinsic incubation period (EIP), the virus is transmissible via the next bite from that mosquito to a susceptible human [20]. Vector competence – the ability of a mosquito to become infected with and ultimately transmit the virus – and EIP are both temperature dependent where higher temperatures in general shorten the EIP and temperature affects transmission in a non-linear fashion [11, 20, 21]. Similarly, mosquito life traits are temperature dependent, and in combination with viral-vector kinetics define the transmission potential of the virus-vector pairing [11, 18, 22-24].

*Ae. aegypti* are known to stay relatively near or within domiciles, and transmission of arboviruses is believed to be moved among households by people, rather than by inter-household mosquito-based transmission [25]. That means that when a household outbreak occurs, it likely began with an index case within the household rather than the introduction of an infectious mosquito, and mosquitoes already within the household become exposed from that human infectious case and then spread to other members of the household [25]. This is supported by evidence that the more time spent at home increased the likelihood of being bitten by *Ae. aegypti* in Iquitos, Peru [26]. Heterogeneity in the biting habits of *Aedes aegypti* mosquitoes has been observed [5, 6, 27] but modeling frameworks do not always explicitly account for the potential for individual-level *Ae. aegypti* heterogeneity in biting.

Biting is commonly represented in model structures such as compartmental or SEIR models as a constant mosquito biting rate, which describes a time-between-bites process that is exponentially distributed. This governs the movement of mosquitoes from susceptible to exposed as well as the movement of humans from susceptible to exposed classes [28-30]. In two individual-based models (IBM), biting was represented as both an activity (0/1) and associated with altered survival (due to risk of death during biting) [31, 32]. Still other efforts have addressed heterogeneity in *Ae. aegypti* biting of humans, highlighting the importance of such for understanding transmission dynamics of arboviruses [15, 27, 33-35].

Through a combination of experimental laboratory work and a computational framework, we investigated the hypothesis that heterogeneity in individual biting habits of *Ae. aegypti* was quantifiable and had the potential to affect small-scale DENV transmission at the individual level.

## Methods

### Determination of Individual Biting Habits

*Aedes aegypti* (Rockefeller colony) were vacuum hatched and placed in a rearing pan with deionized water and fish food as in [18, 36]. The aquatic stages were held at a constant temperature of 28°C in environmental chambers with a photoperiod of 16:8 light:dark hours [18]. On the day after emergence, adults were cold anesthetized, and females were placed in 4-ounce, white disposable paper cartons with screen fastened around the top to provide containment. Cartons were placed in 24°C, 28°C, or 32°C environmental chambers with 16 individual females per temperature group. Females that died within the 2 days immediately following transfer to the 4 oz carton were censored, as death was likely due to handling (n=3). Female *Ae. aegypti* will take a blood meal in the absence of water to satisfy thirst which may artificially inflate biting [37, 38], so mosquitoes were maintained on 10% sucrose solution at all times (i.e., not starved prior to blood-meal offering). Mosquitoes had constant access to oviposition paper which was kept damp throughout.

Beginning on day 2 post-emergence, mosquitoes were offered a 20-minute blood meal consisting of bovine blood in Alsever’s anticoagulant via Hemotek feeding device, daily at sunrise for 23 days (ending on day 25 post-emergence) [39]. *Ae. aegypti* were previously determined to bite repeatedly with highest frequency at 24-hour intervals [40]. The Hemotek arms were threaded through a port in the environmental chamber so that mosquitoes were never removed from the chamber, and thus temperature remained consistent. Cartons were blown on to introduce CO_2_ cues and then the discs were placed directly on top of the screen at the time of feeding. Blood feeding was recorded at each blood meal offering when the presence of fresh (bright red) blood in the abdomen was observed. Only one person observed each feed to eliminate that as a source of variation. Deaths were documented as they occurred. Two biological replicates at each temperature were performed (n=10-16). There were no significant differences in the distributions of measures of interest (total # bites, time to first bite, time between first and second bites, and time between first and last bites) between replicates across all three temperatures (Kolmogorov-Smirnov test), so replicates were collapsed into three temperature groups (Supplemental Information Figures **S1-4**).

### IBM Model Structure and Parameterization

To investigate the role of mosquito biting heterogeneity on small-scale, household transmission, an individual-based model was developed that simulated household transmission patterns using the heterogeneity in biting behavior observed in our experiments. A schematic of the model is given in Supplemental Figure **S5**. Briefly, a household of two human individuals was assumed, one of whom is initially infectious (the index case), with a single susceptible mosquito which is allowed to bite according to her experimentally-determined bite profile. The mosquito may bite each person with equal probability, regardless of infection status. If the mosquito bites the index case, she is exposed until completion of the EIP, at which time she is infectious. The additional human is initially susceptible until bitten by an infectious mosquito. Data from the field suggests that, while mosquitoes bite multiple times in a single gonotrophic cycle, each biting event was from a single source [7]. The simulation stopped once a secondary case was achieved or at the end of 25 days, whichever came first. We also assumed that contact with an infected human resulted in perfect transmission to the mosquito and, likewise, that a bite from an infectious mosquito always resulted in transmission to the susceptible human in the main model. However, we also explored the sensitivity of the model to variation in the probability of transmission from the infectious index case to the mosquito (*β*_v_) as well as the probability of transmission from the exposed mosquito to susceptible humans (*β*_h_).

Model parameters are given in **Table 1**. But briefly, the DENV infectious period in humans was assumed to 8 days [41]. We varied the mosquito biting behavior in separate simulations to match the different mosquitoes in our experiment. The model was realized for 1000 simulations per single mosquito, with its biting parameterized according to its distinct profile. The mosquito bit with a probability of 0 or 1 determined by the experimental data. That is, if a particular mosquito from the data bites on days 2 and 15 from the experimental data, she will only bite on days 2 and 15 of the simulation. The temperature-dependent EIP for DENV virus was defined as 11.5, 7.9, and 6.4 days for 24°C, 28°C, and 32°C, respectively [24].

**Table 1:** Parameter values for the models were either generated in the Christofferson laboratory or taken from the literature.

| Temp | Parameter | Value (rounded up to nearest day) |
| --- | --- | --- |
| 24 | EIP [20] | 12 days |
| 28 |  | 8 days |
| 32 |  | 7 days |
| All | Infectious Period of Human [41] | 8 days |

All individuals (humans and mosquitoes) are initialized at day 1 of the simulation, which was run for 25 days, and the total number of transmission events to the mosquito and secondary transmission events to the susceptible human among all simulations was recorded. This was done to control for variability in timing and to enable direct comparison across mosquito individuals and their biting profiles. The probability of secondary transmission was calculated as the number of simulations where the susceptible human was infected divided by the total number of simulations.

The output from these models was used to determine which aspects of the bite profiles were most correlated with transmission using the “rcorr” function in R. Specifically, correlation matrices were produced to determine the association between the total number of bites a mosquito performed, time to first bite, the total number of times the mosquito became infected out of 1000 simulations, and the total number of times out of 1000 simulations that the susceptible human became infected.

### Local Sensitivity Analysis

A sensitivity analysis was conducted for the human infectious period (+/-2 days) and the EIP (+/-2 days), as well as *β*_v_ and *β*_h_ (0.25-0.75) [36, 42], for each temperature, compared to the parameter values presented in Table 1. The same metrics were calculated and observed for changes from the base scenario.

### Analytic Computation of Mosquito Transmission

We can analytically derive mathematical expressions for the probability that the mosquito is successfully exposed (p_ME_) and the probability that transmission to the second human occurs (p_HE_):

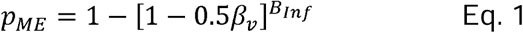

Where B_Inf_ is the total number of bites that a mosquito takes during the human infectious period, i.e. the total number of bites taken during the first 8 days. This expression can be viewed as 1 – the probability that every bite taken during the human infectious period did not result in an exposure.

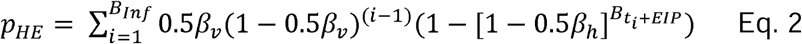

The probability p_HE_ sums over the number of bites taken during the human infectious period. The first terms represent the probability that bite i results in a successful mosquito exposure (0.5*β*_*v*_) multiplied by the probability that previous bites have not yet resulted in exposure, 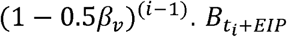 is the total number of bites that a mosquito takes after t_i_+EIP days, where t_i_ represents the day that bite i occurred.

Further, we can calculate the total possible number of transmissible bites by counting pairs of bites separated by the EIP, regardless of where they occur. This gives a measure of transmission potential for each mosquito under relaxed assumptions about synchrony with the index case and is a vectorial capacity-like metric. These computational measures were completed for each mosquito biting profile.

### Exploring common mathematical representation of biting

Commonly used compartmental models assume that the time-between-bites for a given population is distributed exponentially with a single rate parameter, *λ*. We tested this assumption by fitting the empirical data to a null model of exponentially distributed time-between-bites. We estimated the *λ*_TEMP_ parameter of the exponential distribution from the empirical data using the maximum likelihood estimate (sample mean time between bites)^−1^. Because the experimental data were measured discretely (daily), we discretized the continuous ∼EXP(*λ*_TEMP_) to an analogous geometric distribution. The resulting probability distribution was then tested against the time-between-bites of the experimental data using the Chi-square test for goodness-of-fit (*chisq*.*test* function with simulated p-values) to determine whether we could reject the null hypothesis that the experimental data could be derived from the discretized ∼EXP(*λ*_TEMP_) distribution. Further, we tested the proportion of mosquitoes at each simulated scenario that bit at least once and at least twice using the *prop*.*test* function in R. The empirical and theoretically-derived distributions of time-to-first-bite, time-between-first-and-last bites, and total number of bites were compared with the Chi-square goodness-of-fit as above.

For the time-between-bites data where the Chi-square test for goodness-of-fit rejected the null hypothesis, we assessed whether a negative binomial distribution described the data better. To do this we fit both geometric and negative binomial distributions by maximum likelihood estimation and calculated Akaike Information Criterion (AIC) values for model comparison.

## Results

### Experimentally measured heterogeneity in biting among individual mosquitoes

The experimental data showed that biting is heterogenous among mosquitoes at the individual level across temperature conditions. At 32°C, mosquitoes were less likely to bite overall within the timeline of the study (25 days post-emergence) with only 46.2% biting at least once (12/26), and 34.6% biting twice or more (9/26). At 28°C, all (26/26) mosquitoes bit at least once while 84.6% (22/26) bit twice or more. At 24°C, 77.8% (21/27) of the mosquitoes bit at least once and 59.3% (16/27) bit at least twice. The average time between bites was shortest, the average total number of bites higher, and the average daily number of bites highest at 28°C. This provide further evidence that temperatures like 28°C are optimum for transmission [11]. Summary metrics for each temperature are given in Table 2. Further, the distributions of total number of bites, time to first bite, time between first and second bites, and the time between first and last bites are displayed in Figure 1. We explore the time between bites in detail below.

**Table 2:** Summary metrics of biting profiles at each temperature.

| Temperature | Total # that bit 1+ | Total # bit 2+ | Empirical average of total bites (range) | Empirical average Time between Bites | Empirical average daily # bites (range) |
| --- | --- | --- | --- | --- | --- |
| 24 | 21/27<br>(77.78%) | 16/27<br>(59.3%) | 2 (0-5) | 5.99 | 0.083 (0-0.185) |
| 28 | 26/26<br>(100%) | 22/26<br>(84.6%) | 3.92 (0-12) | 3.41 | 0.163 (0.038-0.346) |
| 32 | 12/26<br>(46.15%) | 9/26<br>(34.6%) | 0.96 (0-3) | 5.62 | 0.040 (0-0.154) |

**Figure 1:**
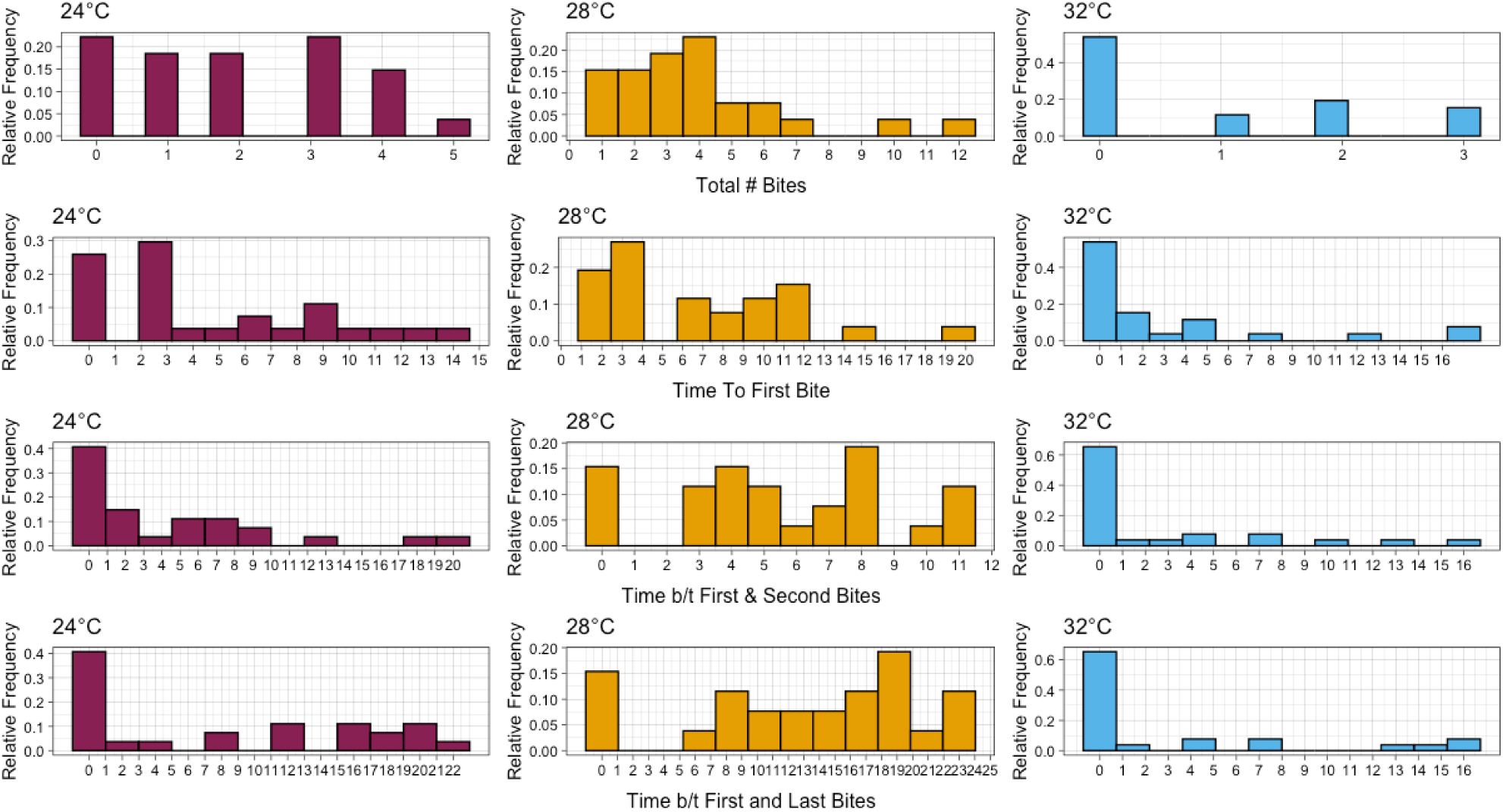
Summary of Ae. aegypti biting characteristics at three temperatures: Total number of bites, time to first bite, time between the first and second bites, and time between first and last bites.

We measured the contribution of individual bite profiles to the likelihood that 1) a mosquito becomes exposed and infectious and 2) a secondary (human) infection occurs given the introduction of a single infectious index case. Not all mosquitoes that bit became exposed and/or transmitted (**Figure 2**). That is, they either did not bite soon enough to become exposed by the viremic index case and/or there were no bites after the EIP resulting in subsequent transmission to susceptible household members. At 28°C, nine mosquitoes that bit at least once never became exposed; at 24°C, 7 mosquitoes that bit did not become exposed; and 3 mosquitoes that bit at least once at 32°C did not become exposed (**Figure 2A.1-C.1**). On the other hand, all the exposed mosquitoes who bit at least twice (6/6) subsequently transmitted with non-zero probability at 32°C. At 24°C, 80% of exposed mosquitoes that bit at least twice (8/10) transmitted to susceptible individuals; and at 28°C 93.75% (15/16) exposed mosquitoes that bite twice transmitted to susceptible household members with a non-zero probability as one mosquito’s only subsequent bite was not outside the EIP (**Figure 2A.2-C.2**).

**Figure 2:**
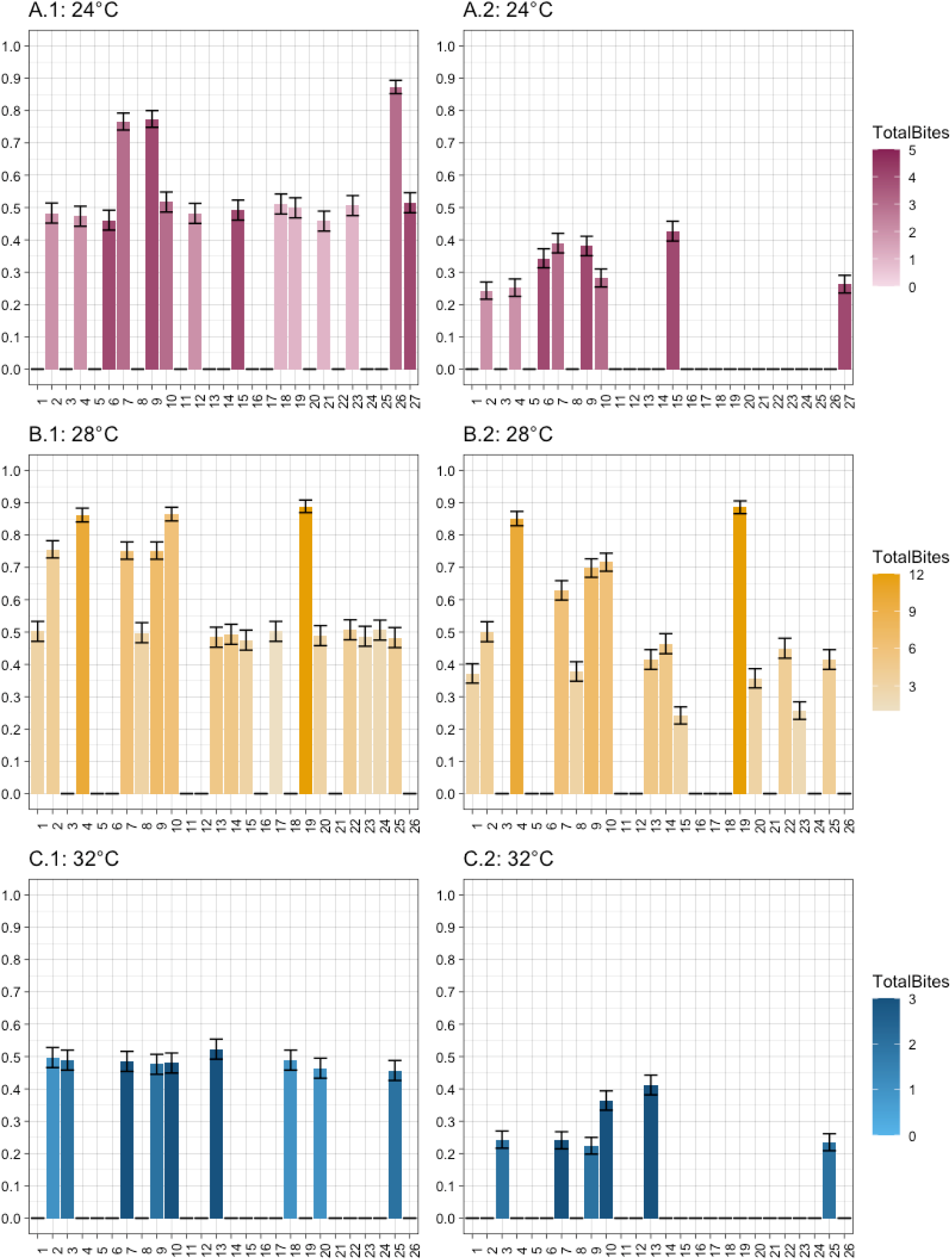
The proportion of mosquito-specific simulations (y-axes) that resulted in successful mosquito exposure from the index case (A-C.1) and subsequent transmission to the susceptible human (A-C.2) at each temperature: 24°C (A), 28°C (B), and 32°C (C). Color gradients indicate total number of bites per individual that transmitted.

To determine whether these differences in patterns were related to the static parameters of EIP and human infectious period, we tested the sensitivity of the model to these quantities. We found that the system was relatively insensitive to small changes in either parameter. The system was not overly sensitive to changes in EIP, which may be an artifact of initializing the model in sync with the index case. Overall, changes in probability were modest (Supplemental Figure S7). At 24°C there was no change in the overall outcome of transmission for any individual mosquito. At 28°C, only two mosquitoes (#23 and #24) had any change in the overall binary outcomes of supporting any transmission or not. With an increase in EIP, #23 did not subsequently transmit after exposure while a decrease in EIP allowed #24 to transmit to the susceptible human. At 32°C, a lengthening of the EIP by 2 days resulted in #3 and #7 having a zero probability of transmitting to the susceptible household member (**Supplemental Figure S7**).

The length of infectious period had more of an impact, likely due to the synchrony of initialization of the model. Nevertheless, at 24°C, a 2-day shorter infectious period changed the overall yes/no outcomes of mosquitoes #10 and 23 to a zero probability of mosquito exposure and subsequent transmission to a susceptible human. (**Supplemental Figure S8**). On the other hand, increasing the infectious period by 1 or 2 days resulted in 4 additional mosquitoes (#8, #3, #22, and #25) having successful mosquito exposure. Only two (#13 and #25) also had subsequent transmission to a susceptible human with a 2-day increase in the infectious period. At 28°C, three mosquitoes (#11, #12, and #26) went from no transmission to successful mosquito exposure with +2 days on the infectious period and two (#11 and #12) also resulted in subsequent transmission to a susceptible human. A shortening of the infectious period resulted in #8, #17 and #25 having no simulations successfully expose the mosquito. Finally, at 32°C, a shorter infectious period resulted in #9 not successfully becoming exposed or subsequently transmitting, while a 2-day longer infectious period resulted in #26 having a non-zero (and relatively high) probability of exposure (∼90%), though this did not result in subsequent mosquito-to-human transmission (**Supplemental Figure S8**). Finally, we varied both the probability of transmission from the index case to the mosquito (Bv) and the probability of transmission from exposed mosquitoes to susceptible humans (Bh) to determine how this parameter altered the overall outcomes. The change in probabilities was largely proportional and did not result in a change in overall yes/no outcomes for any individuals across all three temperatures (**Supplemental Figure S9-10**).

### Correlation of biting profile with simulated transmission potential

The correlation between mosquito exposure and subsequent transmission was uniformly positive, and moderate to high (0.6 at 24°C, 0.90 at 28°C, and 0.73 at 32°C). The total number of bites was highly and significantly correlated with mosquito exposure and subsequent transmission across all temperatures (**Table 3**). Time-to-first bite (TTFB) was also significantly correlated with mosquito exposure across all temperatures but was not significantly correlated at 32°C for transmission to a susceptible human (**Table 3, Supplemental Figure S6**). These relationships were inverse, indicating that a shorter time-to-first bite correlates with higher probability of transmission. While this is likely at least partially due to the model structure, previous work has also suggested that biting sooner is an important component of transmission [18].

**Table 3:** The significantly correlated variables associated with successful exposure/ transmission demonstrate that factors associated with successful transmission are not uniform across all temperatures.

| Event | Temp | Variable | Rho |
| --- | --- | --- | --- |
| Mosquito exposure<br>(IC -> Ms) | 24°C | Time to 1<sup>st</sup> bite | -0.86 |
|  |  | Total # Bites | 0.38 |
|  | 28°C | Time to 1<sup>st</sup> bite | -0.84 |
|  |  | Total # Bites | 0.66 |
|  |  | Time b/t 1<sup>st</sup> and last bites | 0.72 |
|  | 32°C | Time to 1<sup>st</sup> bite | -0.94 |
|  |  | Total # Bites | 0.66 |
|  |  | Time b/t 1<sup>st</sup> and last bites | 0.83 |
|  |  | Time b/t 1<sup>st</sup> and 2<sup>nd</sup> bites | 0.70 |
| Mosquito to Susceptible<br>Human Transmission<br>(Me->Hs) | 24°C | Time to 1<sup>st</sup> bite | -0.61 |
|  |  | Total # Bites | 0.59 |
|  |  | Time b/t 1<sup>st</sup> and last bites | 0.72 |
|  | 28°C | Time to 1<sup>st</sup> bite | -0.75 |
|  |  | Total # Bites | 0.84 |
|  |  | Time b/t 1<sup>st</sup> and last bites | 0.86 |
|  | 32°C | Total # Bites | 0.74 |
|  |  | Time b/t 1<sup>st</sup> and last bites | 0.86 |

The time between first and second bites was not significantly correlated with either transmission event at either 24°C or 28°C but was positively correlated with mosquito exposure at 32°C (*ρ*=0.60). However, the time between first and last bites was significantly, positively correlated with transmission to susceptible household members from exposed mosquitoes across all temperatures. However, only at 28°C and 32°C was this significantly correlated with the initial mosquito exposure itself (**Table 3, Supplemental Figure S6**).

We determined which bites most contributed to transmission out of all successful transmission simulations for both IC->Ms and Me->Hs. To show the hypothetical role of each bite within an individual biting profile, we calculated the proportion of transmission events attributable to a particular bite out of all simulations that resulted in a transmission event (not all bites) (Figure 3). In most cases, the first bite did account for the majority of that mosquito’s transmission when exposure was limited to a single index case as in our model, especially at 32°C where 100% of mosquito exposure occurred upon the first bite. At 28°C and 24°C, there was more variability in the bite at which a mosquito was exposed, though the first bite still accounted for more than other bites.

**Figure 3:**
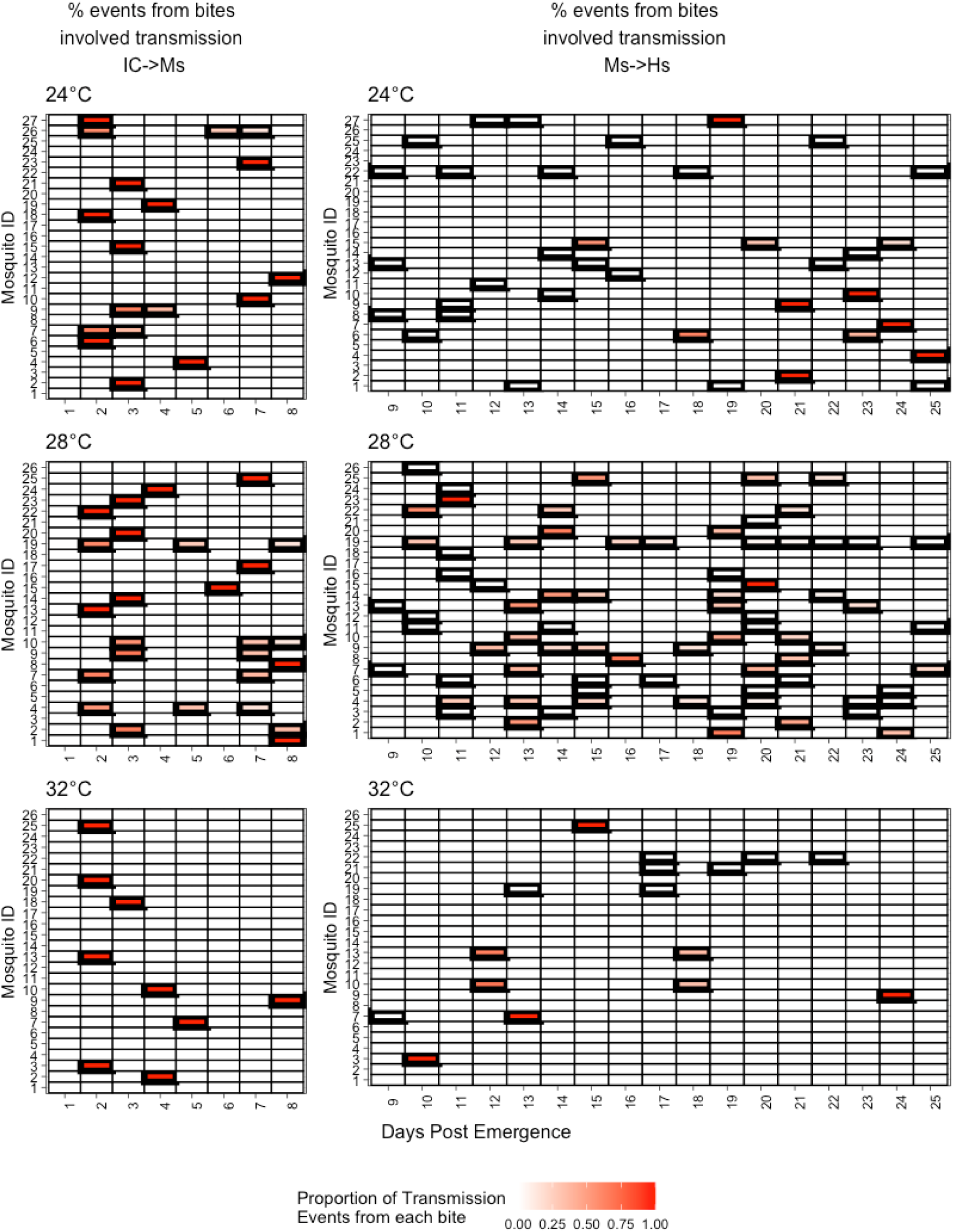
Experimental bite data from mosquitoes shows heterogeneity among individuals. Bolded cells represent days on which a mosquito bit. **Left Column:** During the first 8 days of simulation, gradations of red represent the proportion of IC->Ms events established by that bite out of total bites resulting in IC-Ms transmission. **Right column:** For days 9-25, gradations of red represent the proportion of exposed mosquito to susceptible human events established by that bite out of total bites resulting in Me->Hs transmission.

#### Analytic computation of mosquito transmission

The probability that the mosquito is successfully exposed (p_ME_) and the probability that transmission to the second human occurs (p_HE_) can be analytically derived as above (Eq. 1-2). With perfect transmission (*β*_*V*_ = 1), the p_ME_ is entirely determined by the number of bites that an individual takes during the human infectious period. And once the number of bites taken during the human infectious period is accounted for, p_HE_ is largely determined by how many bites occur EIP days after a successful mosquito exposure. The outcomes of the simulations (Figure 2) are in agreement with these calculations (Supplemental Figure S11). In parallel, we counted the pairs of bites that occurred at least one EIP apart, which relaxes the assumption of asynchrony. This is because it quantifies, for a given profile, how many pairs out of the total bite-pair combinations could theoretically support transmission. This measure is an approximation of the vectorial capacity equation, which is an extension of the Ross-Macdonald framework [15, 43]. Figure 4 demonstrates that at 28°C more mosquitoes have a greater likelihood of transmission due to the timing of pairs of bites based on the temperature-dependent EIP. At 24°C and 32°C, especially, there are fewer overall opportunities to transmit given the timing of pairs of bites, while at 28°C there is much more variability and overall opportunity (Figure 4). This demonstrates that even when the assumption of synchrony between the introduction of the infectious index case and mosquito bite timeline, the potential for consequences on transmission directly related to bite heterogeneity exist.

**Figure 4:**
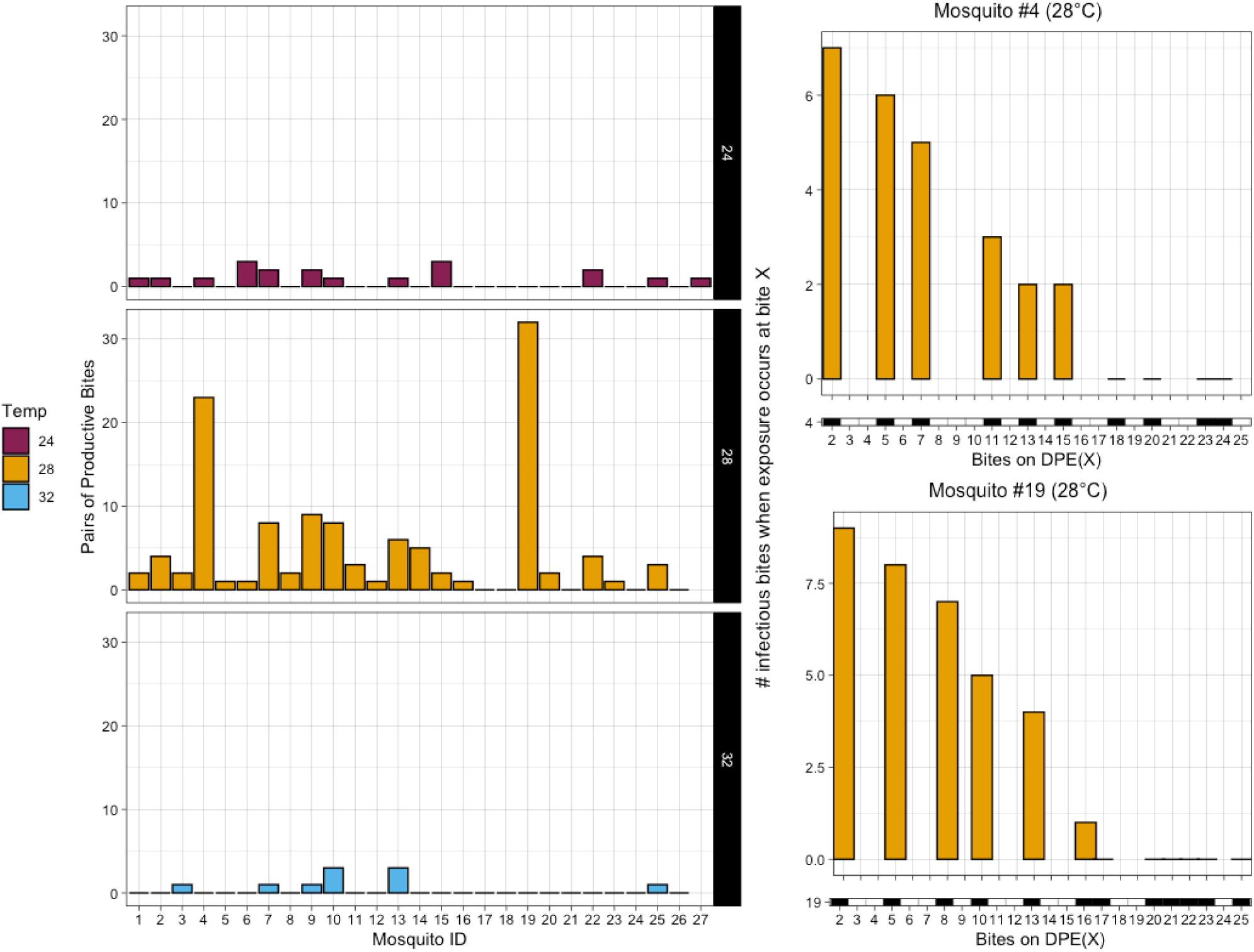
Left: At each temperature, the number of pairs of bites for each mosquito (x-axis) that are separated by at minimum the temperature dependent EIP. Right: Two representative mosquitoes from 28°C with high frequency of biting demonstrating the number of bites occurring after the temperature dependent EIP (histogram) corresponding to each bite (black and white bars below).

### Null model assumptions about biting do not represent the observed process

We investigated whether the distribution of time-between-bites is consistent with the commonly assumed distribution: ∼EXP(*λ*_Temp_) after calculating *λ*_Temp_ as the reciprocal of the sample mean of time between bites (0.167 for 24°C, 0.293 for 28°C, and 0.178 for 32°C). We compared the experimental and theoretical times-between-bites and found that there was a significant difference at 28°C, but not at 24°C or 32°C (**Figure 5, row 1**). We then simulated 1000 hypothetical bite profiles at each temperature using *λ*_Temp_ and compared the proportion of mosquitoes that bit at least once and the proportion that bit at least twice between the experimental and simulated mosquito populations. There was no difference between the empirical data and theoretical population at 28°C in the proportion that bit at least once while both the 24°C and 32°C empirical data had significantly lower proportions (**Table S1)**. When we compared the proportion that bit at least twice, the empirical data had significantly lower proportions compared to the simulated population at all temperatures (**Table S2**).

**Figure 5:**
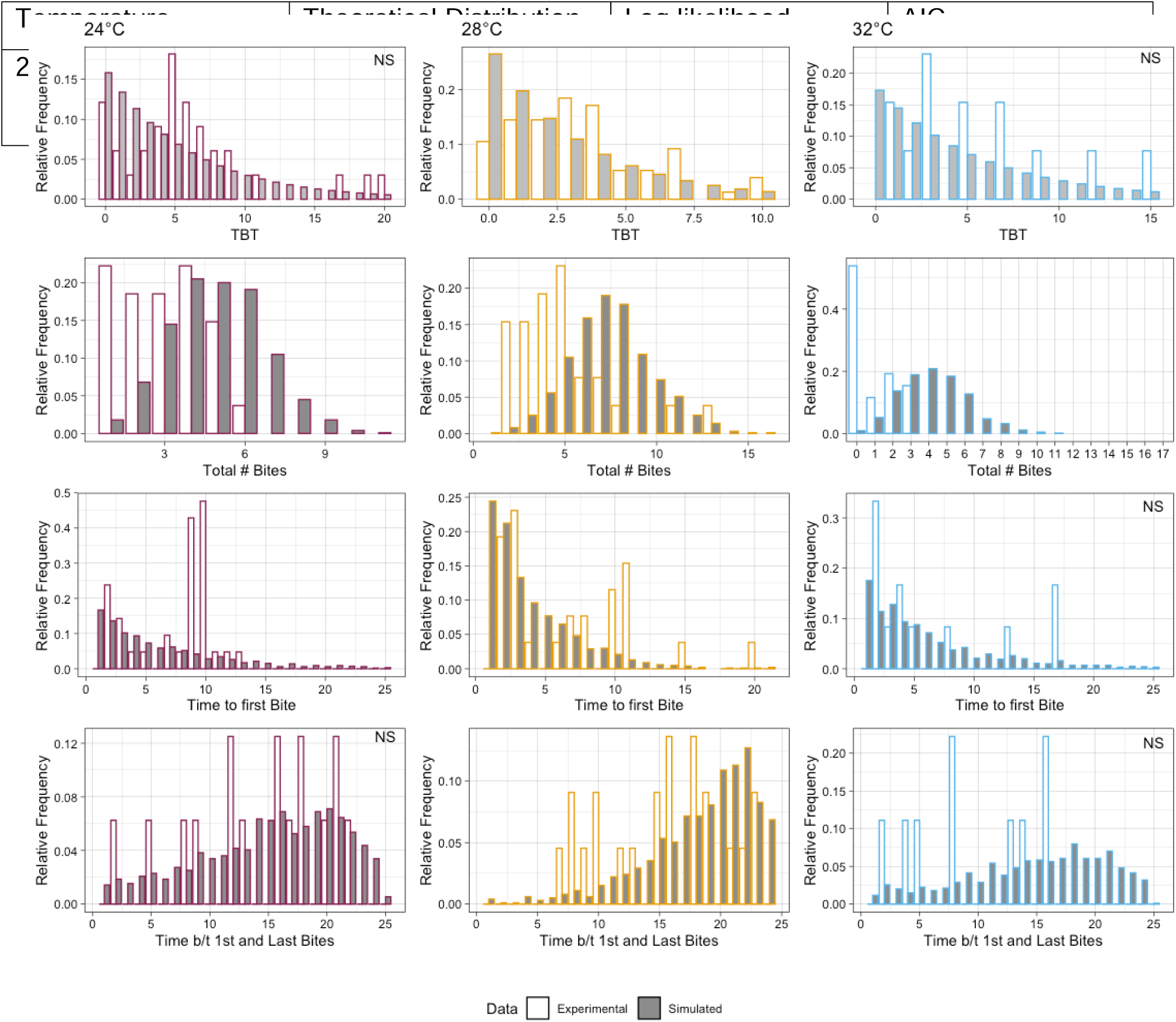
Distributions of metrics describing biting from the experimental data (white bars) to data simulated under the null assumptions of ∼EXP(*λ*_Temp_) (grey bars). Top Row: Time between bites. Second Row: the total number of bites. Third Row: the time to first bite. Bottom Row: Time between first and last bites. Distribution differences were tested usng the Chi-square goodness of fit and NS indicates not significant.

We then compared the distributions of several key metrics found to be correlated with transmission (Table 3). At all temperatures, the distribution of the total number of bites was different between the empirical and simulated populations (**Figure 5, row 2**). The time to first bite was not significantly different for 32°C, again due to overall lack of biting. This metric was significantly different for 24°C and 28°C (**Figure 5, row 3)**. Finally, at only 28°C was the time between first and last bites significantly different, though again the distributions at 24°C and 32°C are visually different and this is again likely due to an overall relative dearth of biting at these temperatures. For 28 degrees, we also assessed whether an alternative discrete distribution with additional parameter, the negative binomial, fit the time-between-bites data better. The negative binomial distribution has a much lower AIC score (difference of -16.79) than the geometric distribution (analogous to a discretized exponential distribution), which provides strong support that this 2-parameter distribution describes the data better than the single parameter geometric distribution.

**Table 4:** Comparison of distributions shows that negative binomial is a better fit to the 28°C bite data (time between bites) compared to the geometric.

## Discussion

The impact of using individual-level data rather than population averages (such as for compartmental models) remains to be elucidated, but it is important to note that the simple act of biting twice does not translate to a mosquito becoming part of a transmission cycle because the timing of bites is important [18]. *Ae. aegypti* biting in the field was estimated as 0.63-0.76 bloodmeals per day (a population estimate) within a single gonotrophic period, while our data suggests this may be lower for some temperatures (Table 2) [7]. Further, different descriptors of the timing of bites were significantly correlated with transmission for each temperature, indicating that a more macro-level heterogeneity across temperature exists as well. However, at this level, it is important to note that biting in the wild and ultimately DENV transmission is affected by a multitude of environmental factors [15, 33, 44].

When we perturbed the model by varying several parameters, the most sensitive variable was the infectious period of the index case, which is likely attributable to the synchrony of initialization. Subsequent exploration of the relative timing of pairs of bites (Figure 5) displayed the number of potential secondary bites quantified, and with this an estimate of an individual vectorial capacity-like quantity can be determined to quantify transmission potential [15, 43]. Further, this shows that supports previous findings that earlier bites are more likely to contribute to transmission [18]. The general lack of sensitivity of the system to changes in EIP is also due to the timing of pairs of bites in the empirical data relative to the length of the EIP, while varying reductions in both βv and βh translated to expected reductions in the probability of transmission. But again, we observed biting in isolation while there is certainly an interplay between mortality, biting, and other factors [15, 35, 43]. In addition, the IBM herein provides a framework with which to test the interaction of biting with these other parameters.

While assessing the goodness of fit of the ∼EXP(*λ*_TEMP_), we determined that at the optimum temperature of 28°C, this distribution did not adequately describe the time between bites. Nor did the temporal measures of mosquito bite profiles simulated from ∼EXP(*λ*_28°C_) represent the observed empirical distribution of total bites, the distribution of time to first bites, or time between the first and last bites (all of which were significantly correlated with successful transmission outcomes). We show that a negative binomial distribution better fits the time between bites at 28°C, suggesting that models describing optimum conditions should consider this alternative fit, or a similar continuous distribution such as a gamma distribution. At 24°C and 32°C, while the ∼EXP(*λ*_TEMP_) could not be ruled out as an appropriate distribution, there are clear quantifiable differences observable and likely this is due to the low number of overall bites compared to 28°C.

We recognize caveats of the study. First, laboratory settings represent ideal conditions for mosquitoes, and partial and full bloodmeals are probably a conservative estimate of mosquito-human contact, as probing can lead to transmission [36]. However, recently published data demonstrate that while there are likely differences in biting habits across geographic and laboratory populations, a degree of heterogeneity was observed in a field-derived colony, indicating that heterogeneity among individuals is not limited to laboratory colonies [36]. Constant temperatures may not capture the nuances of temperature effects, though have been successfully used for important modeling studies and are important to isolate thermal effects [11, 24]. In addition, mortality is differential among temperatures, as demonstrated in several previous studies with obvious implications for transmission and biting [11, 22, 24]. While our study was not designed to explicitly measure mortality, we did find that the mortality of our three temperatures did follow the same pattern as previously published, with 28°C having greater average survival (>25 days) than either 24° or 32° days (24 and 23.5 days, respectively) [11, 24]. Finally, it is important to emphasize that our data are not intended to represent the entire range of heterogeneity potential among individuals, but to explore the implications of possible heterogenous biting profiles on theoretical transmission scenarios.

These data indicate that there is considerable heterogeneity in not only total number of bites, but the timing of bites as well. Further, these metrics were differentially affected by temperature. Several studies have demonstrated the non-monotonic nature of mosquito life traits and/or that the optimal temperature for biting is approximately 28°C [11, 17, 24]. Our data further confirms a thermal optimum for *Ae. aegypti-*driven arbovirus transmission, as well as suggests that the commonly used representation of biting at this optimal temperature may not best represent the heterogeneity and associated arbovirus transmission potential.

## Supporting information

Supplemental Inforation

## Acknowledgements

Thanks to Elizabeth Barrett Browning for the convenient title pun and to Mr. John A. Carriere, Jr. (RCC’s dad) for construction of a custom mosquito-feeding stand and to the reviewers for their helpful feedback.

## Funding

This work was supported by NIH/NIGMS grant R01GM122077 (RCC) and the Burroughs Wellcome Fund 2019 Collaborative Research Travel Grant (RCC). The funders had no role in study design, data collection and analysis, decision to publish, or preparation of the manuscript.

## Supplemental Tables & Figures

**Supplemental Figure S1:** There was no significant difference (Kolmogorov-Smirnov test) in the total number of bites at any of the tested temperatures

**Supplemental Figure S2:** We also saw no significant difference in the distribution of time to first bite across the three temperatures (Kolmogorov-Smirnov test).

**Supplemental Figure S3:** Similarly, there was no significant difference between biological replicates in the time between first and second bites across all three temperatures (Kolmogorov-Smirnov test).

**Supplemental Figure S4:** Finally, there was no significant difference between biological replicates in the time between first and last bites across all three temperatures (Kolmogorov-Smirnov test).

**Supplemental Figure S5**: A schematic of a hypothetical mosquito with an individual bite profile where a bite occurs when M{bite} = 1. Infectious contact from the index case (P_H0_) at Time = 2 results in the mosquito becoming exposed (M{status} = 1). A bite from the M{status} = 1 mosquito at Time = 5 did not result in transmission to a susceptible individual (S_H_) as the EIP had not concluded. However, after the EIP, the mosquito status changes to infectious (M{status} = 2) and bites from the mosquito, such as Time = 12 to S_H_ results in a transmission event and changes the status to S_H_ as “unavailable for infection”.

**Supplemental Figure S6**. Complete correlation matrices for all three temperatures. X indicate p is not significantly different from 0.

**Supplemental Figure S7: Local sensitivity analysis for the extrinsic incubation period of DENV in the mosquito vector:** The proportion of simulations (y-axis) where a particular mosquito (x-axis) becomes exposed (P(IC—Ms)) or transmits (P(M->H)) per a range of extrinsic incubation periods. Only mosquitoes with at least one scenario resulting in a non-zero probability of transmission are shown.

**Supplemental Figures S8: Local sensitivity analysis for infectious period of the index case:** The proportion of simulations (y-axis) where a particular mosquito (x-axis) becomes exposed (P(IC—Ms)) or transmits (P(M->H)) per a range of infectious periods. Only mosquitoes with at least one scenario resulting in a non-zero probability of transmission are shown.

**Supplemental Figure S9: Local sensitivity analysis for the probability of transmission given contact between a susceptible mosquito and the infectious index case (**P_**v**_): The proportion of simulations (y-axis) where a particular mosquito (x-axis) becomes exposed (P(IC—Ms)) or transmits (P(M->H)) per a range of f3_v_. Only mosquitoes with at least one scenario resulting in a non-zero probability of transmission are shown

**Supplemental Figure S10: Local sensitivity analysis for the probability of transmission given contact between an infectious mosquito and susceptible human (**P_**h**_): The proportion of simulations (y-axis) where a particular mosquito (x-axis) transmits (P(M->H)) per a range of f3_h_. Only mosquitoes with at least one scenario resulting in a non-zero probability of transmission are shown.

**Supplemental Figure S11:** The outcomes of mosquito exposure and subsequent transmission to the susceptible household member derived directly from Equations 1 & 2 (see Methods).

**Supplemental Table S1**: Comparison of the proportion of mosquitoes that bit at least once between the experimental data and simulated population. There was a significant difference at 24°C and 32°C, using prop.test function in R.

**Supplemental Table S2**: Comparison of the proportion of mosquitoes that bit at least twice between the experimental data and simulated population. There was a significant difference at across all temperatures, using prop.test in R.

## Notes

### Competing Interest Statement

The authors have declared no competing interest.

### Summary of Updates

Significant revision to the manuscript, including the analyses and text. Underlying data did not change.

