## Supplemental Inforation for "How do I bite thee? Let me count the ways: Exploring the Implications of Individual Biting Habits of Aedes aegypti for Dengue Transmission"

**Christofferson, et al.**

**Supplemental Information**

**Supplemental Figure S1:** There was no significant difference (Kolmogorov-Smirnov test) in the total number of bites at any of the tested temperatures

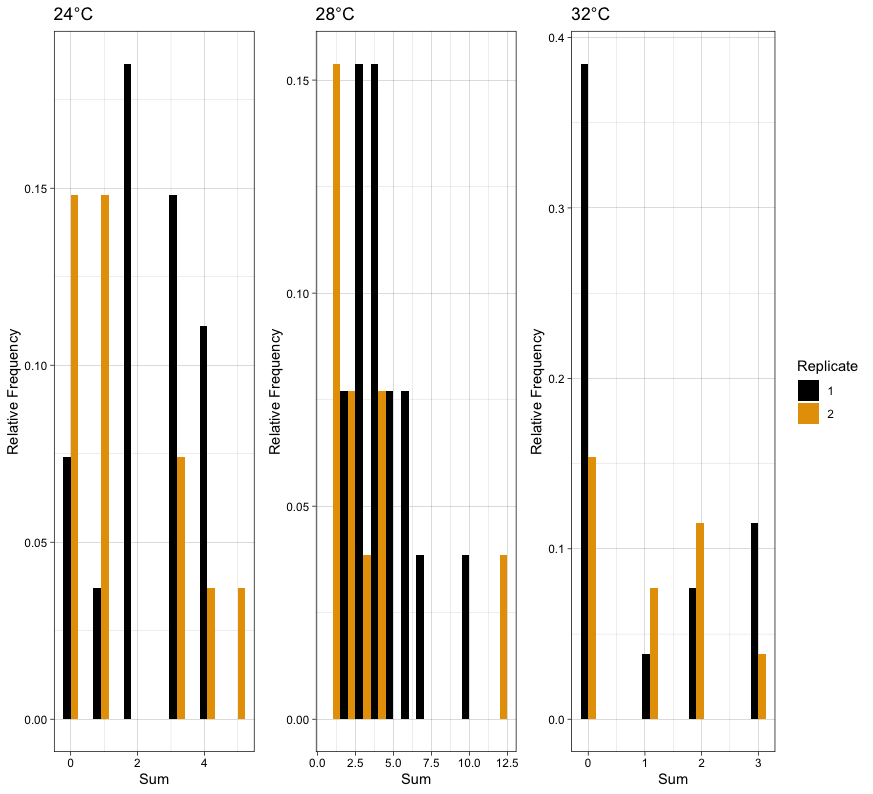

**Supplemental Figure S2:** We also saw no significant difference in the distribution of time to first bite across the three temperatures (Kolmogorov-Smirnov test).

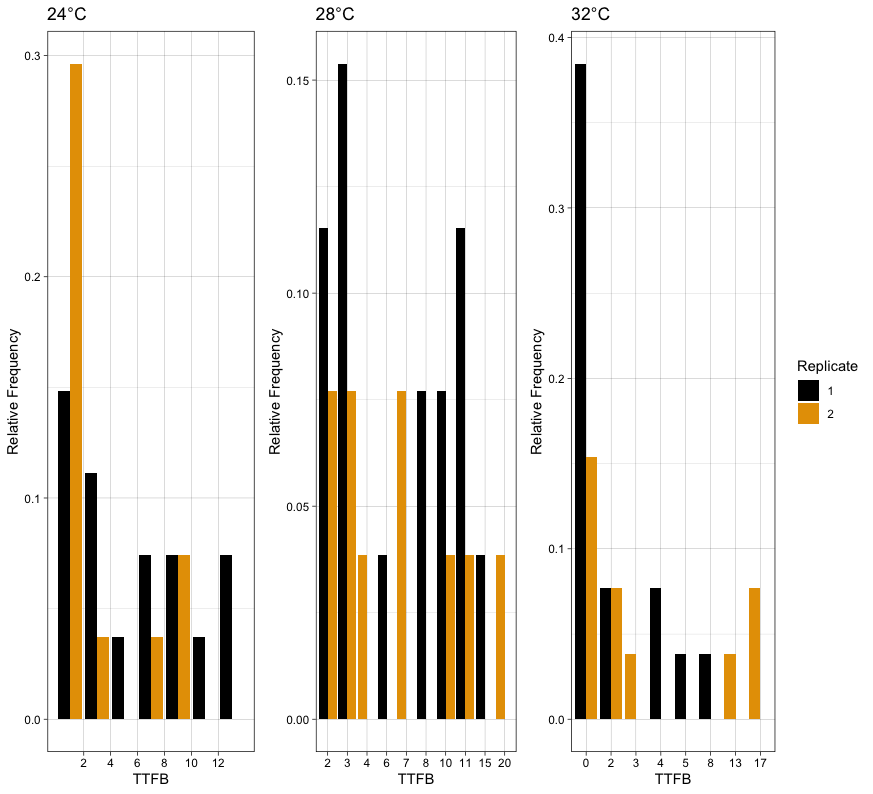

**Supplemental Figure S3:** Similarly, there was no significant difference between biological replicates in the time between first and second bites across all three temperatures (Kolmogorov-Smirnov test).

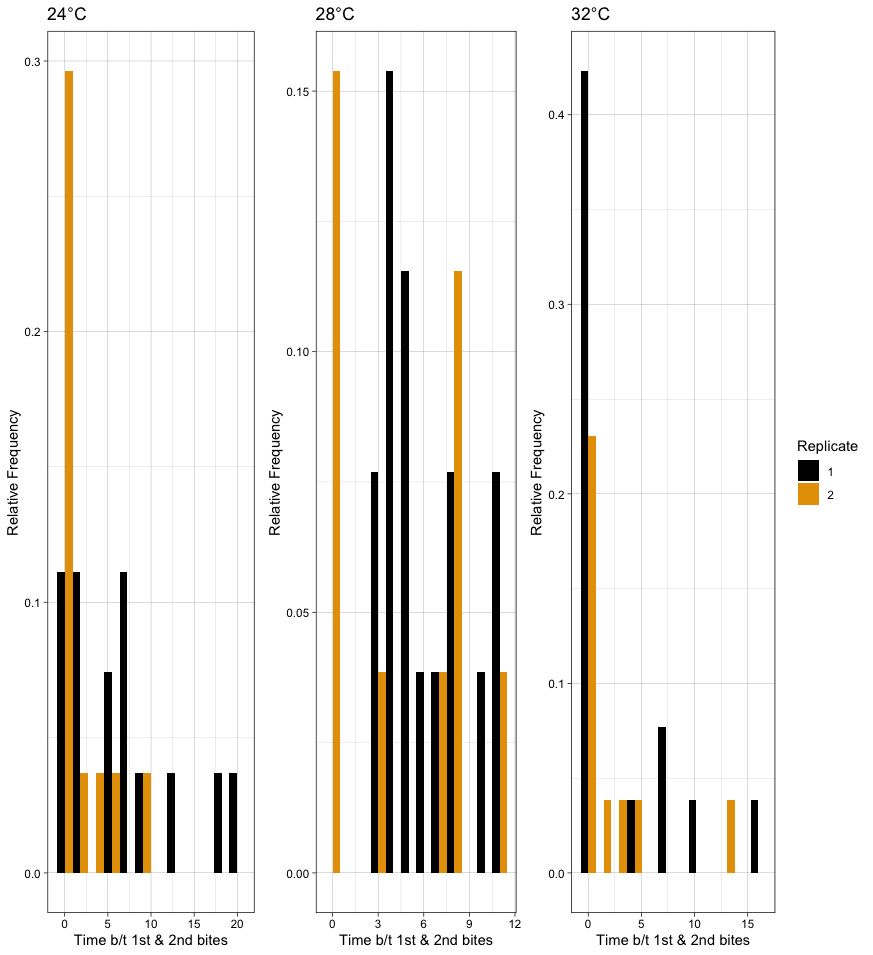

**Supplemental Figure S4:** Finally, there was no significant difference between biological replicates in the time between first and last bites across all three temperatures (Kolmogorov-Smirnov test).
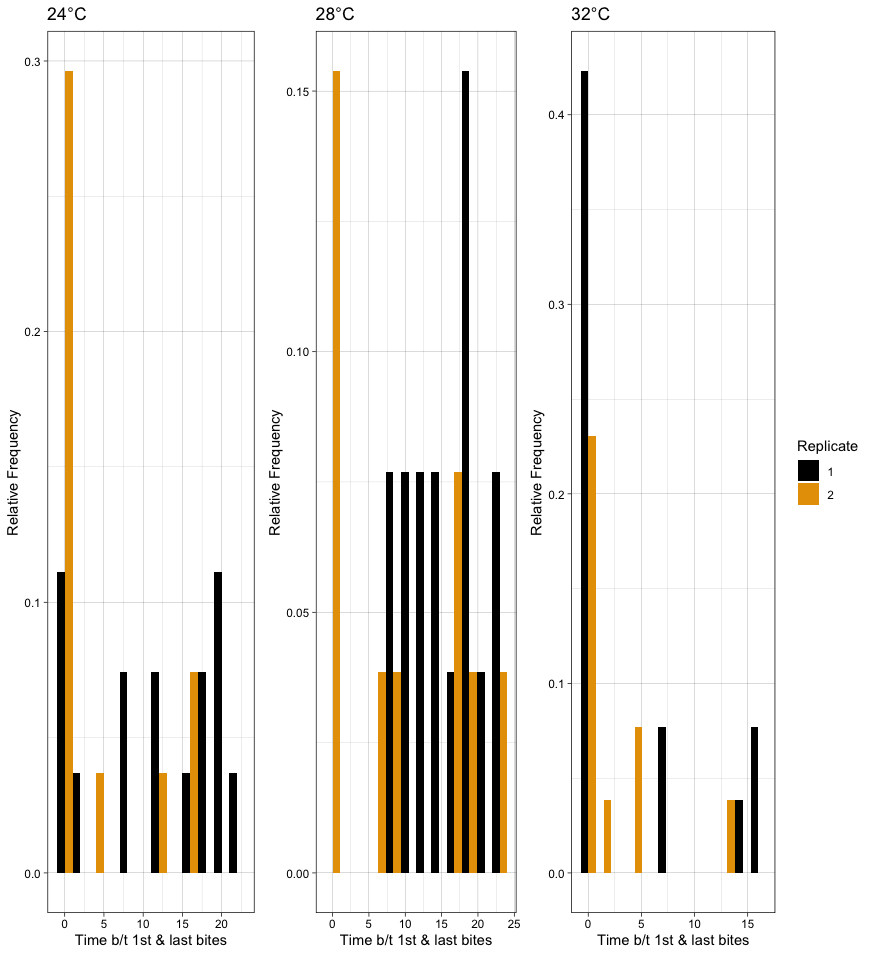

**Supp. Figure S5**: A schematic of a hypothetical mosquito with an individual bite profile where a bite occurs when M{bite} = 1. Infectious contact from the index case (P_H0_) at Time = 2 results in the mosquito becoming exposed (M{status} = 1). A bite from the M{status} = 1 mosquito at Time = 5 did not result in transmission to a susceptible individual (S_H_) as the EIP had not concluded. However, after the EIP, the mosquito status changes to infectious (M{status} = 2) and bites from the mosquito, such as Time = 12 to S_H_ results in a transmission event and changes the status to S_H_ as “unavailable for infection”.

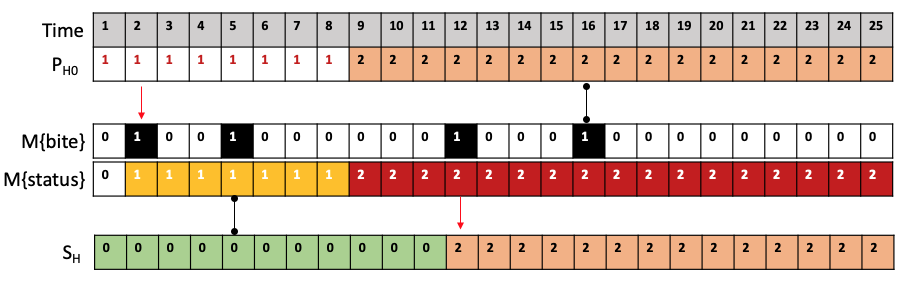

**Supplemental Figure S6.** Complete correlation matrices for all three temperatures. X indicate 𝜌 is not significantly different from 0.

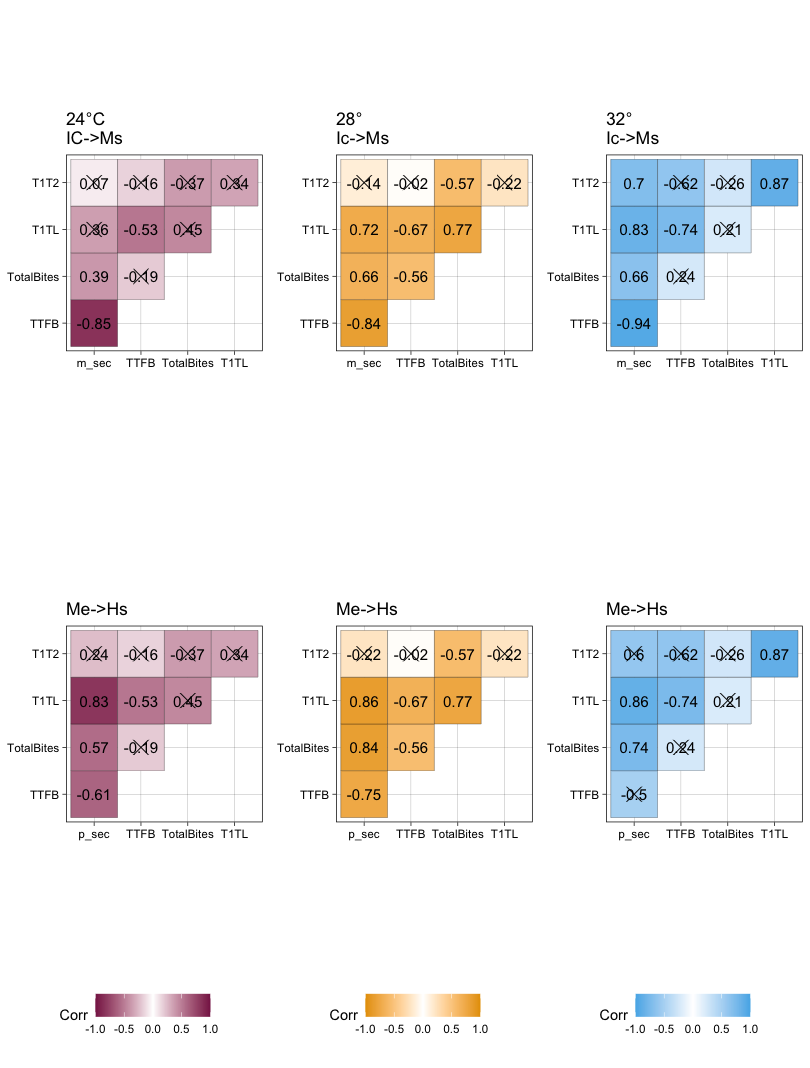

**Supplemental Figure S7: Local sensitivity analysis for the extrinsic incubation period of DENV in the mosquito vector:** The proportion of simulations (y-axis) where a particular mosquito (x-axis) becomes exposed (P(IC—Ms)) or transmits (P(M->H)) per a range of extrinsic incubation periods. Only mosquitoes with at least one scenario resulting in a non-zero probability of transmission are shown.

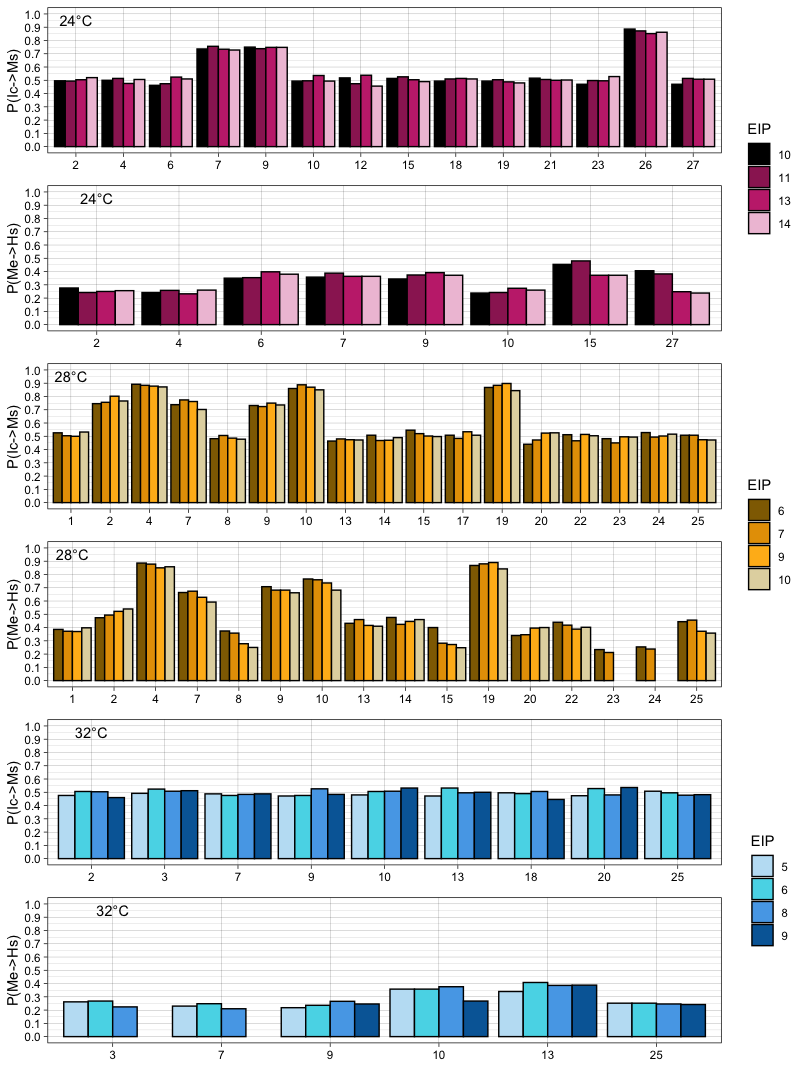

**Supplemental Figures S8: Local sensitivity analysis for infectious period of the index case:** The proportion of simulations (y-axis) where a particular mosquito (x-axis) becomes exposed (P(IC—Ms)) or transmits (P(M->H)) per a range of infectious periods. Only mosquitoes with at least one scenario resulting in a non-zero probability of transmission are shown.

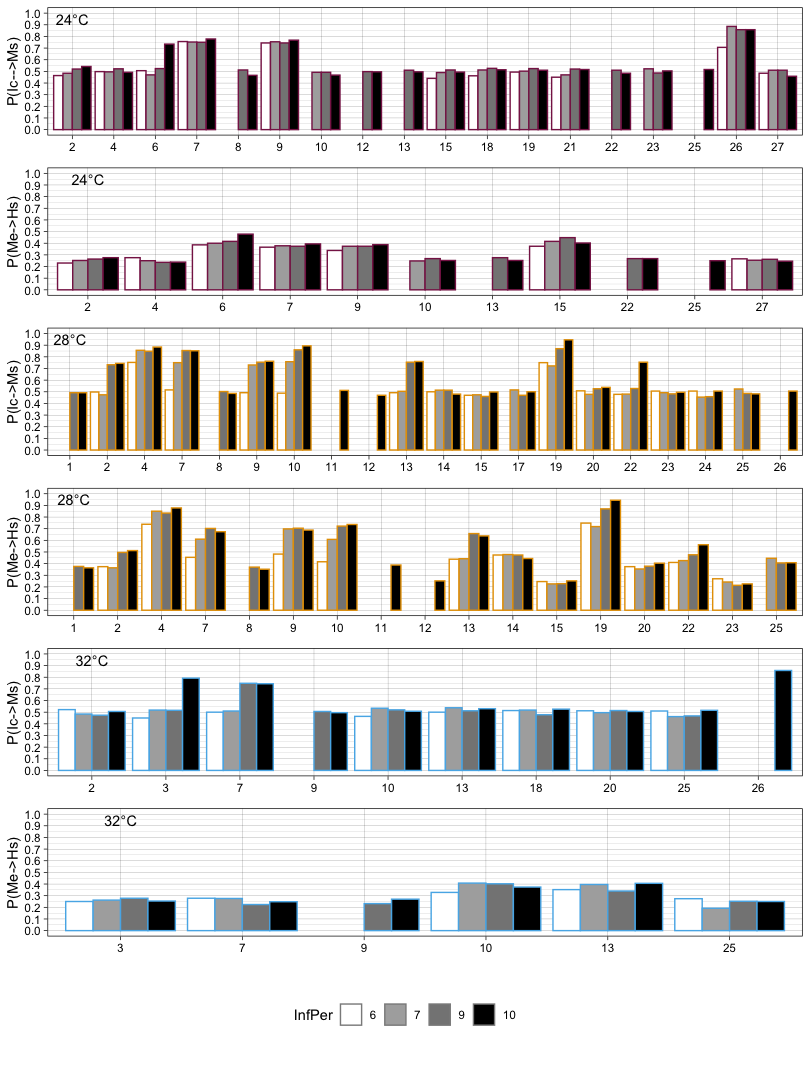

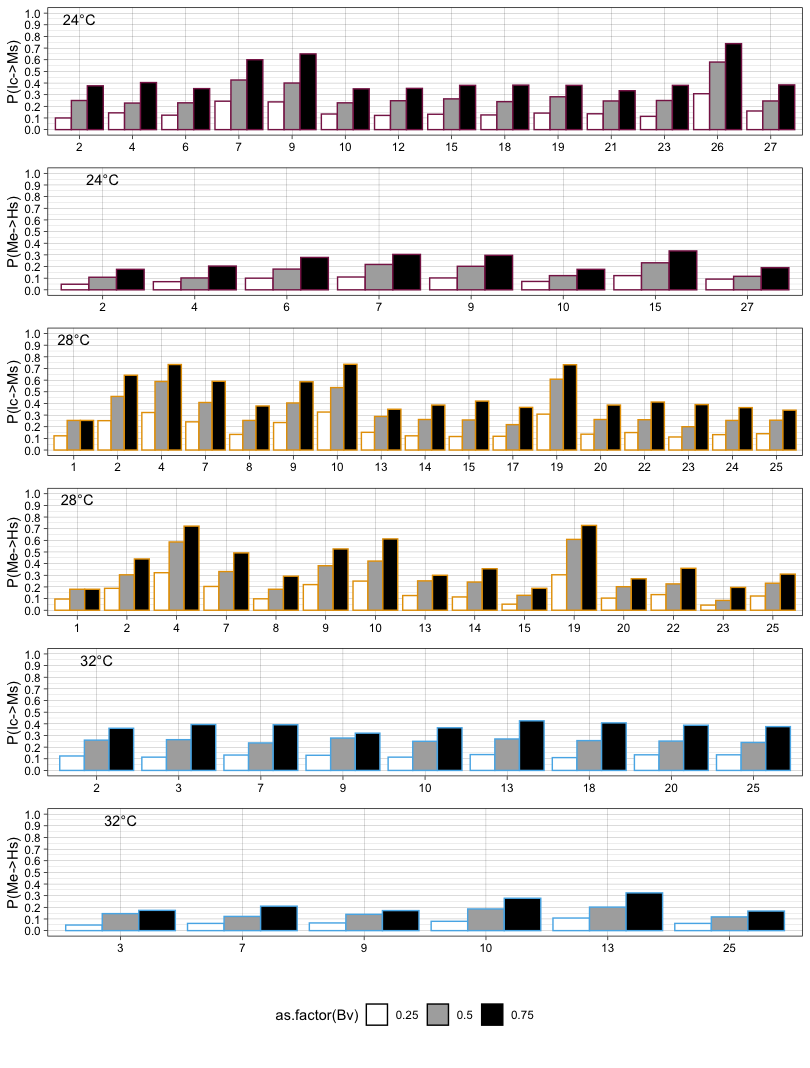
**Supplemental Figure S9: Local sensitivity analysis for the probability of transmission given contact between a susceptible mosquito and the infectious index case (𝛽_v_):** The proportion of simulations (y-axis) where a particular mosquito (x-axis) becomes exposed (P(IC—Ms)) or transmits (P(M->H)) per a range of 𝛽_v_. Only mosquitoes with at least one scenario resulting in a non-zero probability of transmission are shown.

**Supplemental Figure S10: Local sensitivity analysis for the probability of transmission given contact between an infectious mosquito and susceptible human (𝛽_h_):** The proportion of simulations (y-axis) where a particular mosquito (x-axis) transmits (P(M->H)) per a range of 𝛽_h_. Only mosquitoes with at least one scenario resulting in a non-zero probability of transmission are shown.

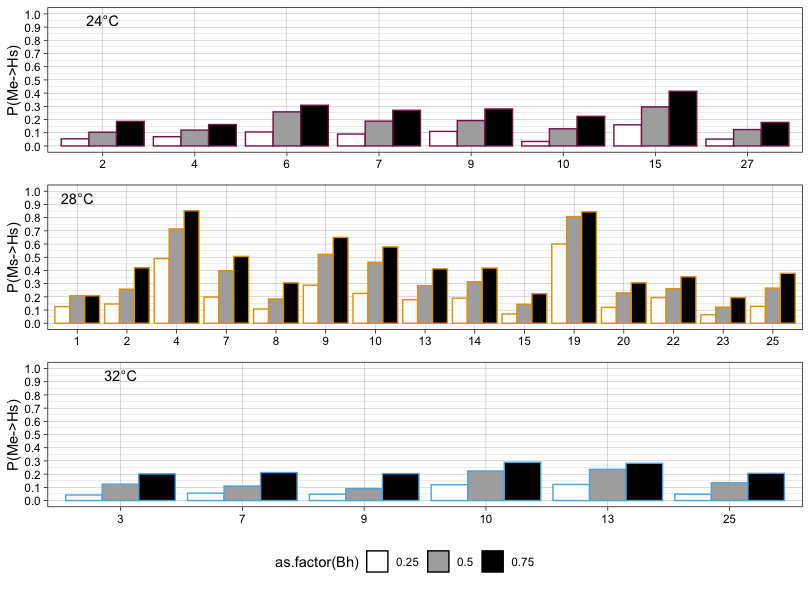

**Supplemental Figure S11:** The outcomes of mosquito exposure and subsequent transmission to the susceptible household member derived directly from Equations 1 & 2 (see Methods).

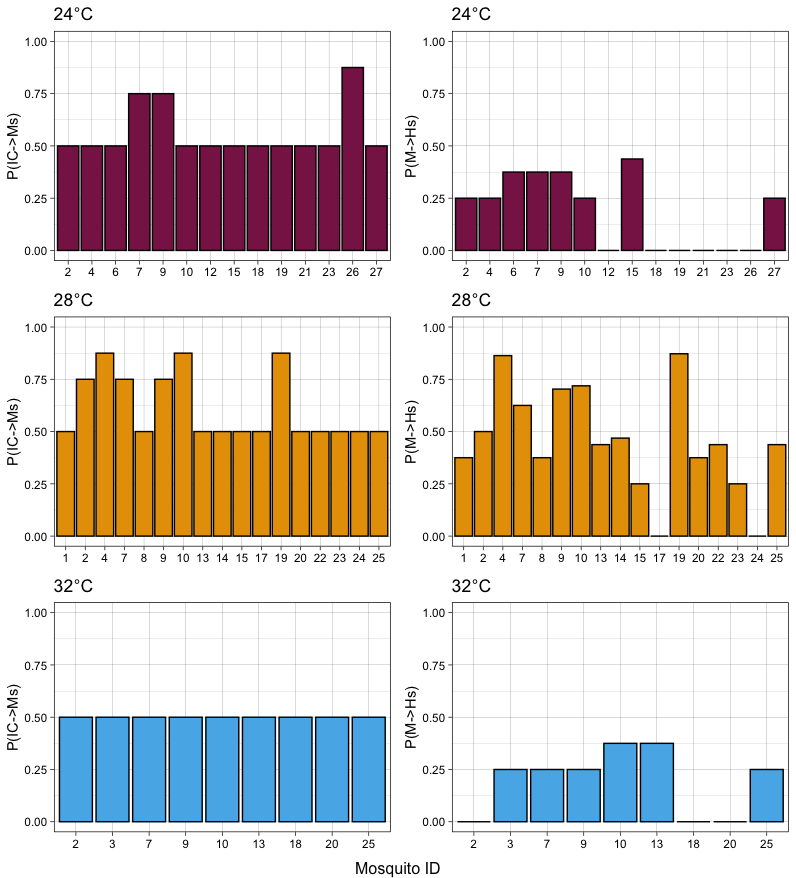

**Supplemental Table S1**: Comparison of the proportion of mosquitoes that bit at least once between the experimental data and simulated population. There was a significant difference at 24°C and 32°C, using prop.test function in R.

| Temperature | Data Type | Proportion |
| --- | --- | --- |
| 24 | Experimental | 0.778 |
|  | Simulated | 0.914 |
| 28 | Experimental | 1 |
|  | Simulated | 1 |
| 32 | Experimental | 0.462 |
|  | Simulated | 0.914 |

**Supplemental Table S2**: Comparison of the proportion of mosquitoes that bit at least twice between the experimental data and simulated population. There was a significant difference at across all temperatures, using prop.test in R.

| Temperature | Data Type | Proportion |
| --- | --- | --- |
| 24 | Experimental | 0.593 |
|  | Simulated | 0.914 |
| 28 | Experimental | 0.846 |
|  | Simulated | 0.991 |
| 32 | Experimental | 0.346 |
|  | Simulated | 0.940 |
